# Self-Supervised Discovery of Discrete Local States in Noisy Image Sequences

**DOI:** 10.64898/2026.09.17.751938

**Authors:** Song-Chuan Zhao, Daisuke Mizuno

## Abstract

Scientific image sequences often feature recurring structures with unknown appearance and dynamics. While traditional methods using predefined templates, noise models, or motion classes work well when targets are known, such assumptions can hinder exploratory analysis where discovering that knowledge is the primary goal. This work presents a self-supervised framework that maps noisy video into a discrete vocabulary of local states and their spatiotemporal relationships. The method operates without clean targets, pretrained representations, semantic labels, feature templates, or prescribed trajectories, assuming only that informative structures recur for a finite duration within a bounded spatial neighborhood. A temporally masked vector-quantized network predicts the missing central frame from its neighbors, while a masked Transformer refines token assignments by weighing encoder evidence against contextual compatibility. Project-specific token sets define the feature family, while a support model quantifies the temporal evidence for individual occurrences, facilitating optional conservative suppression. The framework is first demonstrated on synthetic videos of moving particles. Without access to clean frames or particle coordinates during learning and selection, the learned states recover compact, point-spread-function-like structures. Tests on low-signal particle-tracking benchmarks yield frame-wise component precision between 87.0% and 94.5%, with recall decreasing as particle density rises. An experimental example using *E. coli* further shows that discrete states can separate biological structures from illumination and acquisition artifacts. In all cases, the primary outputs are interpretable state, activity, support, and component maps; rendered images serve as diagnostics rather than optimization targets.

## 1 Introduction

Feature detection is rarely assumption-free. Classical detectors specify responses for edges [1], invariant keypoint methods identify stable local structures [2], and image registration exploits local intensity and motion constraints [3]. Quantitative particle tracking typically begins with a localization model before linking detections into trajectories [4] or applying global constraints on motion and acquisition [5]. While these assumptions are strengths when the target class is known, they can be limiting in exploratory experiments where the form, count, and dynamics of relevant structures are unknown. In such cases, fixing a template or motion law in advance may hide unanticipated states or force diverse observations into a familiar description.

Self-supervised denoising minimizes reliance on priors by predicting withheld measurements from correlated neighbors without needing clean targets. This approach relies on the noise being independent between held-out and supplied measurements, while the underlying signal remains shared. This principle of estimating clean loss from noisy data alone mirrors Stein’s unbiased risk estimate [6]. Noise2Void masks individual pixels and predicts them using the remaining image content, whereas Noise2Self generalizes this approach by ensuring that predictions at withheld coordinates are independent of the values at those same coordinates [7, 8]. Alternatively, Noise2Noise predicts one noisy realization from another independent observation of the same signal [9]. Since both estimate the same conditional mean, masking and pairing serve as two different ways to provide supervision.

Sequence-based methods apply these templates to the temporal dimension. DeepInterpolation adopts the masking template at a frame-level granularity, withholding the complete target frame and predicting it from adjacent raw frames [10]. DeepCAD uses the pairing template, constructing noisy input–target pairs from consecutive, temporally interleaved volumes [11], which assumes these volumes represent the same underlying activity. Both rely heavily on frame-level temporal redundancy. To handle cases where rapid motion makes a target frame poorly predictable, SUPPORT introduces a spatial blind-spot branch that receives the current frame but excludes the target pixel, complementing a temporal branch that processes neighboring frames [12]. This allows same-frame spatial evidence to preserve signals that are missing or displaced in the temporal sequence.

The present framework uses the masking template but differs in its ultimate goal. Rather than focusing on pixel-level restoration, a vector-quantized bottleneck [13] maps recurrent local image structures into a bounded vocabulary, which is kept as the primary data. A separate masked Transformer [14, 15] then uses token identities and learned relative offsets to evaluate compatibility within the spatiotemporal neighborhood. In contrast, SUPPORT combines convolutional features from neighboring frames and a spatially masked current frame to restore pixels. The computational distinction is specific: the implementation performs contextual comparison on a small set of discrete token positions—each summarizing a *u × u* block—instead of extracting relations from a large raw-pixel stack. While this reduces the cost of the contextual step, the encoder and decoder still process pixels, so it does not inherently lower end-to-end runtime. The key contribution is that attention-refined states remain available for occupancy, co-occurrence, transition, temporal-support, and component analysis without requiring a predefined feature template or motion model. Because every later stage reads the same compact token map, downstream analysis is also inexpensive, counting occupancy, co-occurrence, transitions, and support over tokens instead of recomputing them from pixels; this low per-project cost favors training and applying a dedicated model for each dataset rather than relying on one large pretrained model.

These ideas are organized into three phases (Fig. 1). Phase A learns a vocabulary of local morphology. Phase B assigns project-specific feature roles and qualifies individual occurrences. Phase C groups the retained sites into components for object-oriented analysis. Phase A is *domain-free* in the sense that it requires no clean references, pretrained representations, semantic classes, feature templates, or prescribed trajectories. It does, however, rely on architectural choices like the basic unit and temporal window, and assumes that informative structures recur within a bounded displacement. Phases B and C then interpret this vocabulary to reflect the researcher’s scientific interest, without altering the underlying representations learned in Phase A.

**Figure 1:**
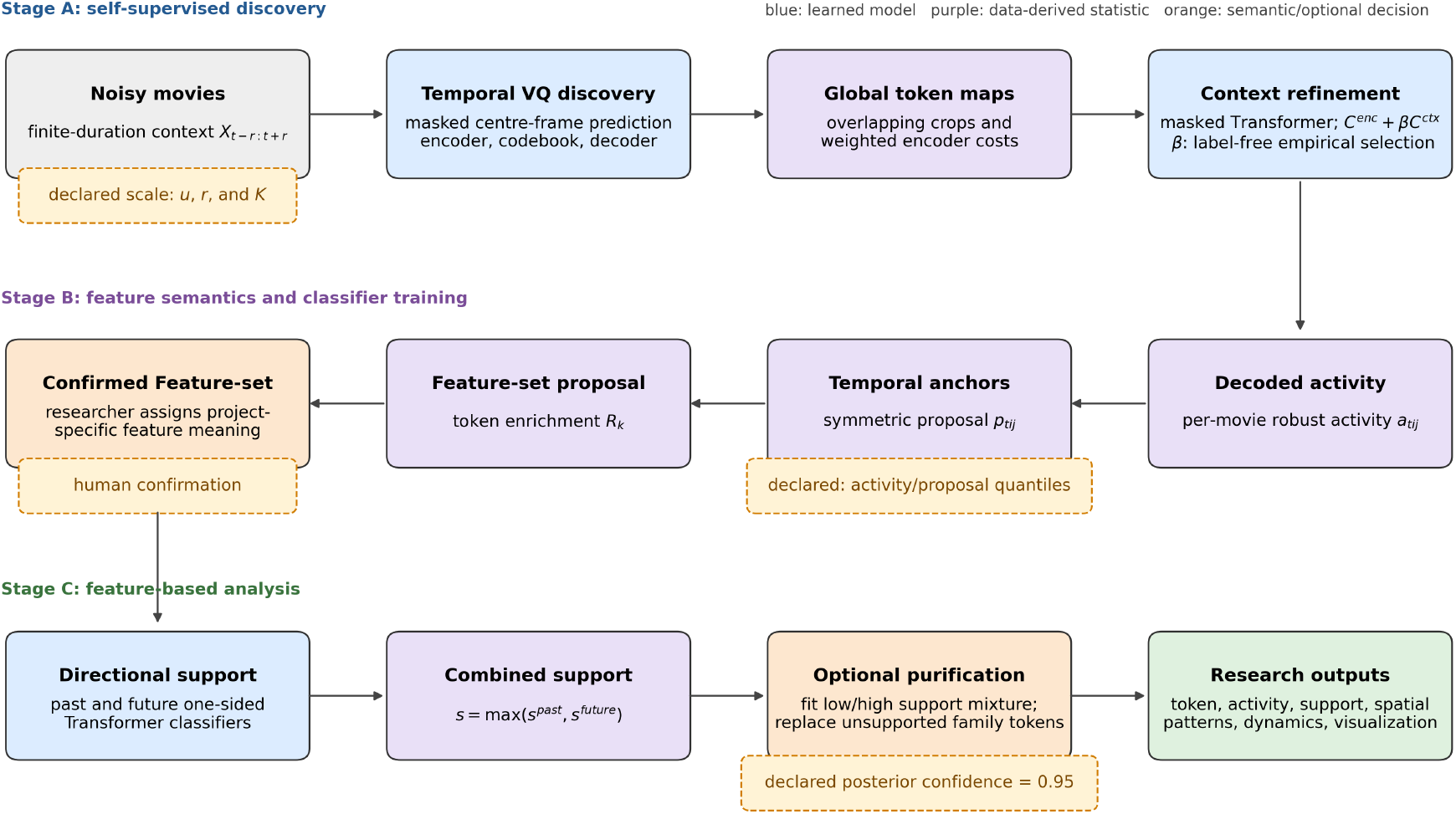
Processing architecture from noisy videos to discrete research outputs. Blue boxes are learned models, purple boxes are statistics derived from their outputs, orange boxes are semantic or optional decisions, and the green box collects quantities available for analysis. Dashed orange labels identify non-universal declared choices. In particular, the activity/proposal quantiles and the unsupported-component confidence are analysis settings; after the confidence is fixed, the support-mixture boundary itself is fitted from the noisy data.

The resulting discrete map serves two primary scientific purposes. First, it allows for the refinement of the feature population through contextual reassignment, directional support, and optional suppression. While restoration provides a visual check, minimum pixel error is not the optimization target; uncertain reconstructions may be discarded to improve population purity. Second, state occurrence, activity, support, contextual compatibility, and component structure provide the data for analysis. Since a physical object may be represented by a compatible sequence or spatial combination of local states rather than a single invariant code, training on the specific collection under investigation is appropriate for a descriptive analysis of that collection.

The following sections describe the framework, demonstrate its utility on synthetic data, and evaluate it using benchmark and experimental videos. The framework is defined in Sec. 2 and demonstrated with synthetic data in Sec. 3. Section 4 presents simulated low-SNR and experimental microscopy test cases, and Sec. 5 discusses design choices and limitations. Run-specific training settings are collected in Appendix A.

## 2 Proposed Framework

The framework decouples representation learning from project-specific interpretation (Fig. 1). In the first stage, it learns a global vocabulary of local states, generates a token map for each video, and estimates contextual compatibility without relying on semantic labels. The second stage defines project-specific feature families, incorporating directional temporal support and optional suppression. The third stage introduces a component definer and characterizes the resulting objects. Because the learned vocabulary remains fixed, researchers can adapt the definition and post-processing of feature families to suit a particular analysis.

### 2.1 Framework Overview and Processing Logic

Consider a noisy video where *X*_*t*_ ∈R^*H×W*^represents frame *t*. A basic spatial unit *u* is defined, which corresponds to the characteristic width (in pixels) of the smallest local feature scale the representation aims to resolve. Other spatial dimensions are derived from a single geometry rule:

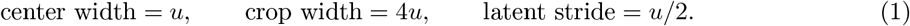

Under this rule, every crop spans 8 *×* 8 latent sites; adjusting *u* scales the total downsampling factor of the encoder and the corresponding mirrored decoder. The model architecture is further defined by the temporal radius *r* and vocabulary size *K*. These are set at the dataset level, as they determine the expected duration of recurrence and the total number of discrete states allowed in the representation, respectively. Their roles are discussed further below.

At each latent-grid location (*i, j*) and time *t*, the framework tracks three distinct quantities: a discrete state *z*_*tij*_, a decoded activity *a*_*tij*_, and—once a feature family is selected—its temporal support *s*_*tij*_. Here, *i* and *j* refer to row and column coordinates on the latent grid rather than image-pixel coordinates. These are maintained as separate values because they address fundamentally different questions: which local pattern was assigned, the strength of its expression, and whether there is compatible temporal evidence. This logic guides the architecture’s progression from a global learned vocabulary to a local, evidence-qualified population of features.

All learned quantities and thresholds discussed here are derived directly from the noisy videos. When available, clean synthetic frames and particle coordinates are reserved exclusively for retrospective evaluation. While training on the full collection is permitted for descriptive analyses of that specific data, held-out sets are used only when claiming generalization or gaining computation efficiency.

### 2.2 Temporally Masked Vector-Quantized Representation Learning

For a crop centered at time *t*, the spatial image content supplied to the encoder is:

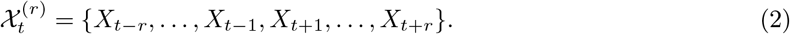

While the noisy central crop *X*_*t*_ serves as the training target, it is excluded as an image channel to maintain the masking objective. Instead, the implementation uses two non-spatial statistics from the full temporal window (including the central crop): the median *l*_*t*_ and a robust scale *q*_*t*_ estimated via the median absolute deviation. Both the target and the context frames are normalized using the same local affine transformation:

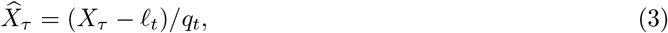

where *q*_*t*_ is subject to a dataset-derived lower bound. In this way, temporal masking strips the spatial pattern of the central frame while preserving its contribution to two permutation-invariant normalization statistics, which are saved to restore local intensity coordinates later. The central frame’s spatial pattern is withheld because severe noise degrades within-frame spatial evidence, whereas structure that recurs across frames survives averaging over independent noise realizations.

Normalized context frames are stacked as channels for a 2D convolutional encoder. Its downsampling blocks consist of strided convolutions and GELU activations, with the number and strides configured to match the latent stride defined in Eq. 1. A final 3 *×* 3 convolution and a hyperbolic tangent activation produce a grid of *d*-dimensional latent vectors *h*_*tij*_ ∈ R^*d*^. The decoder is constructed as a mirror of the encoder, beginning with a 3*×* 3 convolution followed by transposed-convolution upsampling blocks. A codebook of *K* vectors *e*_*k*_ ∈ R^*d*^exists in the same latent space; each encoder vector is quantized via nearest-neighbor assignment:

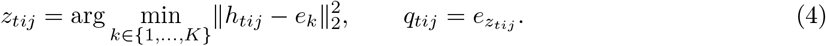

A convolutional decoder then maps this quantized grid back to a prediction of the central crop. To prevent outer pixels from dominating the training process, reconstruction loss is evaluated only within the central *u × u* pixel region, while the rest of the crop provides necessary spatial context. The objective function is:

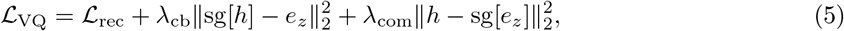

where sg stops the gradient, and *λ*_cb_ and *λ*_com_ weight codebook and commitment losses. Because independent noise varies across frames, it cannot be predicted from neighbors; however, recurrent structures can reduce the expected loss in the center patch. Once training is complete, the encoder and codebook are frozen to ensure that any subsequent changes maintain a stable discrete meaning. The initial decoder is kept as a reference but is not the final renderer. Following contextual reassignment, a copy of the decoder is fitted to the refined, frozen token maps. Since the encoder and codebook remain fixed during this process, the spatial appearance associated with the revised token population is updated without altering the vocabulary itself.

### 2.3 Global Token-Map Assembly

To avoid tying the representation to an arbitrary spatial tiling, inference is performed using overlapping crops (stride=*u/*2 *< u*). This means a single global latent site is typically observed across multiple crops. If crop *n* provides latent vector *h*_*ni*_ for global site *i* (where *i* denotes the flattened grid coordinates), the cost for code *k* is 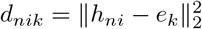. The aggregated encoder cost is then:

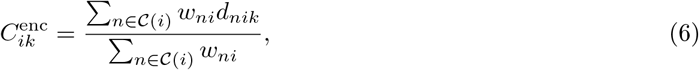

where (*i*) denotes all crops covering the site. A Gaussian weight *w*_*ni*_ is used to decrease the influence of the site as its distance from the crop center increases. Consequently, the crop that places the site closest to its center has the strongest influence, while other overlapping observations serve primarily to mitigate boundary and tiling effects. The initial global token is selected as arg min_*k* 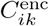_. Rather than keeping only the winning token, all code costs and accumulated weights are retained to ensure that subsequent changes remain traceable to their original encoder evidence.

### 2.4 Context-Guided Token Refinement

Encoder evidence alone cannot determine if neighboring state assignments are mutually compatible. To address this, a masked token predictor is trained on the global token videos. For every eligible site, the central token is removed and a neighborhood with a temporal radius of two (*r* = 2) and a spatial radius of one, resulting in 44 context positions. Token and offset-position embeddings (dimension 32) are passed through a single pre-normalized Transformer encoder layer featuring four attention heads, a 128-dimensional feed-forward sublayer, GELU activation, and 0.1 dropout. During training, context tokens are replaced by a learned dropout-token identity with a probability of 0.15. The masked token is then predicted via mean pooling over context positions, layer normalization, and a linear *K*-class head using a cross-entropy objective. Case-dependent optimization and validation settings are detailed in Appendix A.

Denote the predictor’s negative log-probability of assigning code *k* at site *i* as 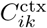. The initial token 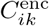 is then refined as follows:

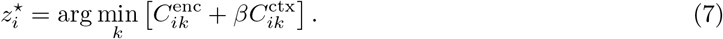

The predictor is trained a single time; *β* is introduced only during the reassignment phase to balance local encoder fit against contextual compatibility. The value of *β* is determined through a fixed, label-free sweep. Let *L*_*β*_ and *D*_*β*_ represent the mean contextual negative log-likelihood and the mean relative encoder cost for a candidate *β*, where candidates range from *β* = 0 to *β*_max_. The following are defined:

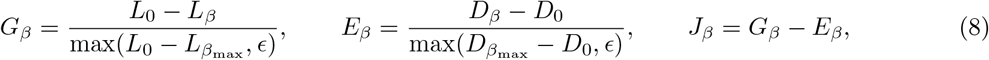

The value that maximizes *J*_*β*_ is then selected. While changed-token fractions are reported for diagnostic purposes, they do not influence this score. After selecting *β*, a separate decoder refinement is performed to fit the frozen, refined tokens to their normalized noisy central crops, while the encoder, codebook, and contextual predictor remain unchanged.

### 2.5 Decoded Activity and Temporal Proposal

The refined decoder maps each token assignment back to a local spatial pattern. Using the saved location and local scale values, these predictions are converted back to video intensity coordinates. Given the dark-background illumination of the data, features are identified by high intensity. If *d*_*tij*_ is the resulting grid-scale intensity and *q*_0.50_, *q*_0.995_ are its per-video quantiles, activity is defined as:

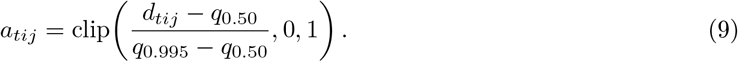

No second site-wise normalization is used at this stage; instead, a single scale is shared within a video to preserve existing temporal and spatial activity differences. Conversely, separate scales for different videos prevent variations in density or illumination from affecting thresholds across the dataset. The impact of inter-video variation is discussed in later sections. Additionally, in cases where the primary challenge is spatial or temporal background inhomogeneity, the decoded intensity can be used without local scales, as shown in Sec. 4.2.

Activity measures expression at one site but does not by itself establish recurrence. The first temporal statistic therefore asks whether a current site is bracketed by compatible activity on both temporal sides within a bounded displacement. This symmetric proposal is used to construct conservative anchor sites for global feature-family discovery. For time separation Δ*t* and spatial displacement (Δ*y*, Δ*x*), define:

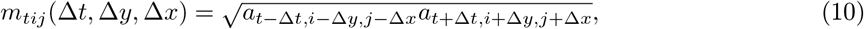

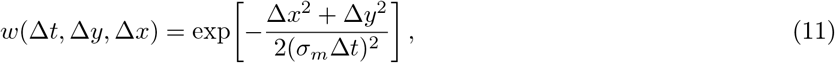

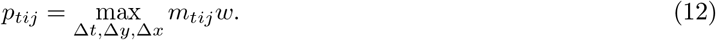

The use of a geometric mean ensures that activity must exist on both temporal sides, while *w* gently favors smaller displacements without imposing a specific trajectory. Contributions from invalid temporal boundaries or paths that cross video blocks are set to zero. A high proposal value *p*_*tij*_ thus indicates a site with strong evidence of temporal consistency; these high-confidence anchors reveal which vocabulary states are enriched at recurrent features. This proposal is deterministic and guides family discovery. Currently, anchor candidates must exceed both the 0.95 activity quantile and the 0.90 proposal quantile within their respective video blocks.

### 2.6 Project-Specific Feature-Family Definition

The vocabulary captures all recurrent structures found across the videos. However, before qualifying local detections, the analysis must identify which codes correspond to the features of interest. These roles are project-specific: a single learned state might be a desired central feature in one study, provide contextual evidence in another, or serve both or neither role. Vocabulary tokens are first ranked based on their representation *efficiency*. For simplicity, the set of feature codes is drawn from this ranked pool; a more structured alternative and its associated component definition are detailed in Sec. 2.9.

First, a broad evidence population is formed from sites that exhibit high activity, high symmetric proposal, and a spatial local maximum of their joint score, as described in Sec. 2.5. For each code *k*, its occupancy within this population is compared against that across all temporally eligible sites. Using additive smoothing, the primary ranking score is:

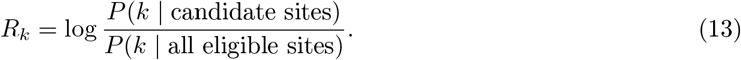

The enrichment *R*_*k*_ quantifies how effectively code *k* concentrates the recurrent, high-activity population relative to its background occupancy. Codes that fail to meet minimum occupancy or evidence count thresholds are excluded. The remaining codes are ordered by *R*_*k*_, with evidence count and mean joint activity–proposal serving as secondary deterministic criteria. This run-specific ranking is recorded, and the top two codes are proposed as the default family for human confirmation. Researchers may revise this selection, as the ideal breadth of the family depends on whether the project targets a single morphology, several related states, or a broader class of recurrent structures.

The *background code* is identified separately from sufficiently occupied states by searching for those with low decoded intensity, activity, and temporal proposal. Low enrichment alone is not a sufficient criterion, as a rare, depleted artifact does not necessarily represent the typical background. Human confirmation simply assigns scientific meaning to these existing states and roles without modifying the vocabulary or the frozen representation models.

### 2.7 Directional Temporal Support

While global feature-family selection answers which codes are relevant, it does not guarantee that every occurrence of those codes is credible. To evaluate individual occurrences, a one-sided support classifier is used that considers activity from only one temporal side within a spatial search neighborhood. The central frame is excluded, and token identities are masked to prevent the classifier from simply replicating the context refinement in Sec. 2.4. This branch projects activity values and relative positions into a latent sequence, which is processed by a Transformer encoder to output a scalar support score trained via binary cross-entropy. This architecture can be applied to either the past or the time-reversed future. Although a single parameter-shared classifier would suffice under time-reversal symmetry, separate models are fitted for each direction. This ensures that irreversible dynamics or acquisition asymmetries—such as illumination dimming—remain observable rather than being forced into a symmetric model. Each branch utilizes a single Transformer layer with 32-dimensional embeddings, four attention heads, 0.1 dropout, and 0.05 context dropout. Case-dependent optimization settings are provided in Appendix A.

While the symmetric proposal of Eq. 12 is used for global family discovery, it is computed separately for each direction when generating classifier labels; this ensures each branch evaluates evidence from a single temporal side rather than the two-sided maximum. Within a given branch, a positive anchor consists of a confirmed-family token with high current activity and high proposal on that branch’s temporal side. Background negatives are non-family tokens with low activity and low one-sided proposal. Family-token negatives include any confirmed-family assignment with low proposal on that branch’s side, including bright assignments that lack temporal evidence in that direction. As a result, the past and future models address distinct directional questions using parallel, rather than bidirectional, label sets. These are pseudo-labels derived from the activity and proposal maps of the same processing pipeline. The support classifier thus acts as an operational refinement of this evidence rather than an independent validation. Excluding the current site and masking token identity ensures the classifier addresses a distinct occurrence-level temporal question.

Both classifiers share the same training/validation frame partition, with an isolated contiguous block withheld from each training video for evaluation. For each direction, the checkpoint with the lowest validation binary cross-entropy is retained. The frozen models are then applied to all processed videos to produce 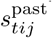and 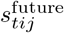, with combined support defined as:

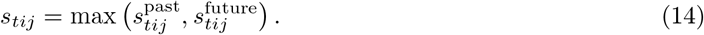

The use of the maximum is intentional. Since a real feature may enter or leave the field of view or be temporarily occluded, valid evidence may not exist on both sides of every frame. Nevertheless, the two directional maps are preserved separately, as their asymmetry serves as a useful temporal statistic.

### 2.8 Support-Guided Purification

A featured token indicates that the local encoder and contextual predictor have selected a member of the chosen vocabulary subset. However, a low support value suggests that the occurrence lacks the temporal evidence central to the framework. Such instances are treated as likely assignment errors induced by noise or context, rather than as weak but valid features. This interpretation justifies an optional, conservative purification step.

The operational cutoff is derived from combined support values at current family-token sites. These values are transformed into logit space and fitted using a two-component Gaussian mixture, where the lower-support component represents the unsupported population. The cutoff is defined as the largest support value for which the posterior probability of membership in this component is at least 0.95. While the confidence level is a preset conservative analysis setting, the mixture parameters and the resulting boundary are fitted directly from the noisy videos. Any featured token at or below this cutoff is replaced by the confirmed background code. This suppression modifies the discrete state map rather than thresholding pixels or optimizing for visual similarity.

In addition to its role in purification, support is maintained as a primary output for use in subsequent stages. When later refinement is applied, the support models and maps are frozen; in this case, retained but weak components may have their activity adjusted based on temporal neighbors, and short enclosed gaps may be filled using compatible token patterns. Maintaining state changes, support evidence, and activity scaling as separate entities allows for a direct comparison between the full learned population and a conservative purified one, while providing an audit trail for each local decision.

### 2.9 Modular Component Construction

Following the steps in Secs. 2.2 through 2.8, the retained feature-family sites are grouped into components, which serve as the operational units for subsequent post-processing. Although this grouping is described late in the pipeline, its definition should be decided in conjunction with the feature-family set in Sec. 2.6. For a simple feature family, the component finder groups sites via eight-connected adjacency, requiring at least one site. The component center in image-space coordinate is determined by the local maximum of the refined-decoder image within the component neighborhood; if this maximum exceeds a permitted deviation from the highest-support site, the coordinate falls back to the center of that latent site. The current design does not fit subpixel coordinates.

A structured alternative optionally establishes two feature sets: a central or core set and a context set. In this approach, a component consists of one central site accompanied by several context sites. The previously defined feature-family set serves as a natural choice for the central set. After freezing the central codes, context codes are estimated independently. Token occurrences within a configurable spatiotemporal neighborhood of accepted central sites are compared against corresponding eligible control occurrences using a smoothed log-enrichment ratio. Spatial offsets are constrained by |Δ*y*| + |Δ*x* | ≤*r*_*c*_; at the default *r*_*c*_ = 1, same-frame context includes the four cardinal neighbors and excludes diagonals. Since the default temporal radius for context selection is zero, context roles are inferred from spatial co-occurrence within the same frame. Codes that exhibit at least two pooled near-center occurrences and positive enrichment are added to the context set. The central and context sets are not necessarily exclusive; a code may label its own site as central while providing context for another center. However, context-only occurrences cannot initiate a component. In the structured component finder, a same-frame candidate must consist of one central site and at least two context sites within its radius-one cross, with overlapping neighborhoods being merged. The merged component’s coordinate is assigned using the same rule as before: the local refined-decoder maximum is used unless it exceeds the permitted deviation from the highest-support central site, in which case the coordinate defaults to the center of that site.

## 3 Framework Demonstration with Synthetic Data

The discovery and purification sequence detailed in Secs. 2.2–2.9 is first tested using simulated point features. The dataset consists of thirty-two videos (128 *×* 128 pixels, 64 frames) with two to ten moving particles per video. Each particle is modeled as a Gaussian point-spread function (PSF) with a 1.5-pixel standard deviation and an amplitude *A* = 1 + *ϵ*, where *ϵ* ~ *N* (0, 0.25^2^) and values below 0.05 are clipped. Consequently, roughly 95% of unclipped amplitudes fall between 0.51 and 1.49, and the saved realization spans 0.381–1.625. Independent Gaussian noise with a standard deviation of 0.5 is added to all pixels. Particles started with random directions and speeds sampled uniformly from 0.15 to 0.75 pixels per frame.

Velocity was perturbed at each frame by independent Gaussian acceleration (*σ* = 0.04 pixels/frame^2^), with speed clipped to the initial interval and trajectories reflected at the image boundaries. This produced persistent yet evolving motion, keeping per-frame displacement small compared to the PSF width. To ensure a fair evaluation, particle positions, clean videos, and the generator PSF are withheld during training, family selection, support fitting, and cutoff determination.

### 3.1 Stage A: VQ Model and Context Refinement

The basic unit is set to *u* = 8 pixels, which, following Eq. 1, results in a 32 *×* 32 pixel crop and a 4-pixel latent stride. Other hyperparameters include a temporal radius of *r* = 2 and a vocabulary of *K* = 16 codes (dimension 32). The encoder thus employs two stride-2 convolutional downsampling blocks with 32 and 64 channels, leading to the 32-channel latent projection detailed in Sec. 2.2. The model is trained on 4096 temporal crops sampled from the noisy videos, predicting the central region from the four surrounding frames.

As described in Sec. 2.3, each frame is assigned a token map which can be used to accumulate an empirical patch for each code. Normalized empirical patches for each initial code are shown in Figure 2. While states 5 and 8 capture compact, bright structures at various sub-patch locations, the most frequently assigned codes describe low-intensity background. This suggests the encoder learned a useful codebook for the subsequent processing stages. We however emphasize that the average patches are empirical and are not the exact representation kernels or basis vectors learned by the model.

**Figure 2:**
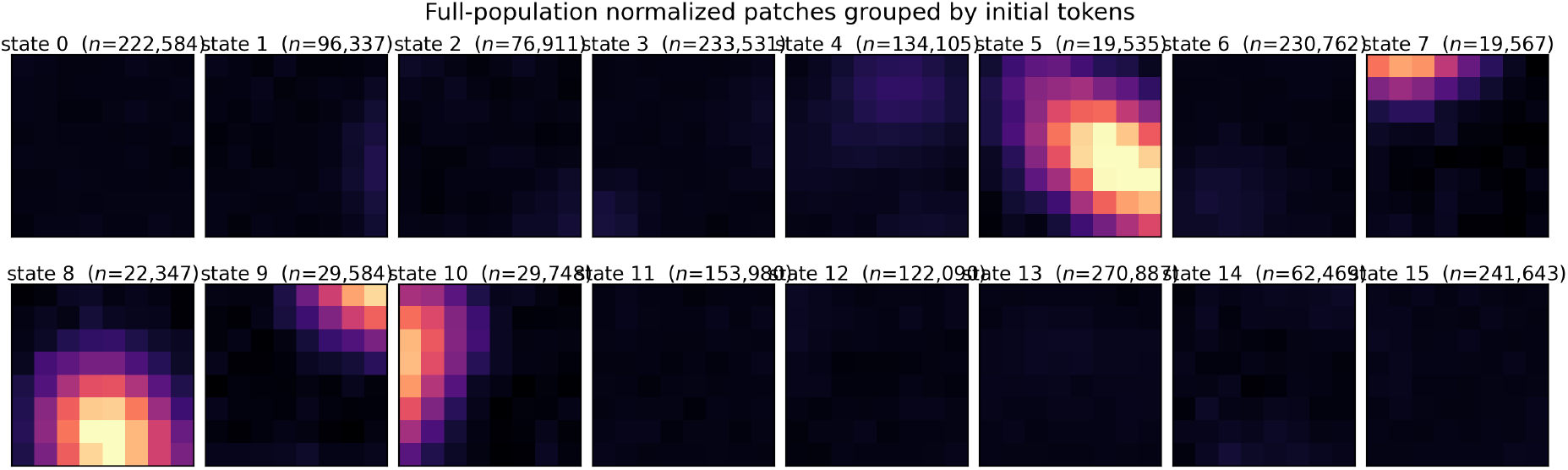
Normalized average patches grouped by the initially learned token assignments. Several states contain a localized bright structure, whereas others describe weak, displaced, or background-like configurations. Counts report assignments in the full token-evidence population.

A one-layer, four-head masked-token Transformer (32-dimensional embedding) is trained using the global state maps from Sec. 2.3. After a single training pass, a sweep is performed over *β* ∈ {0, 0.5, 1, 2, 4, 8} in Eq. 7 without further retraining. Figure 3 shows that while the contextual negative log-likelihood improves as *β* increases, this gain comes at the expense of higher encoder disagreement and a larger fraction of reassigned sites. Based on the label-free score in Eq. 8, *β* = 2 is selected.

**Figure 3:**
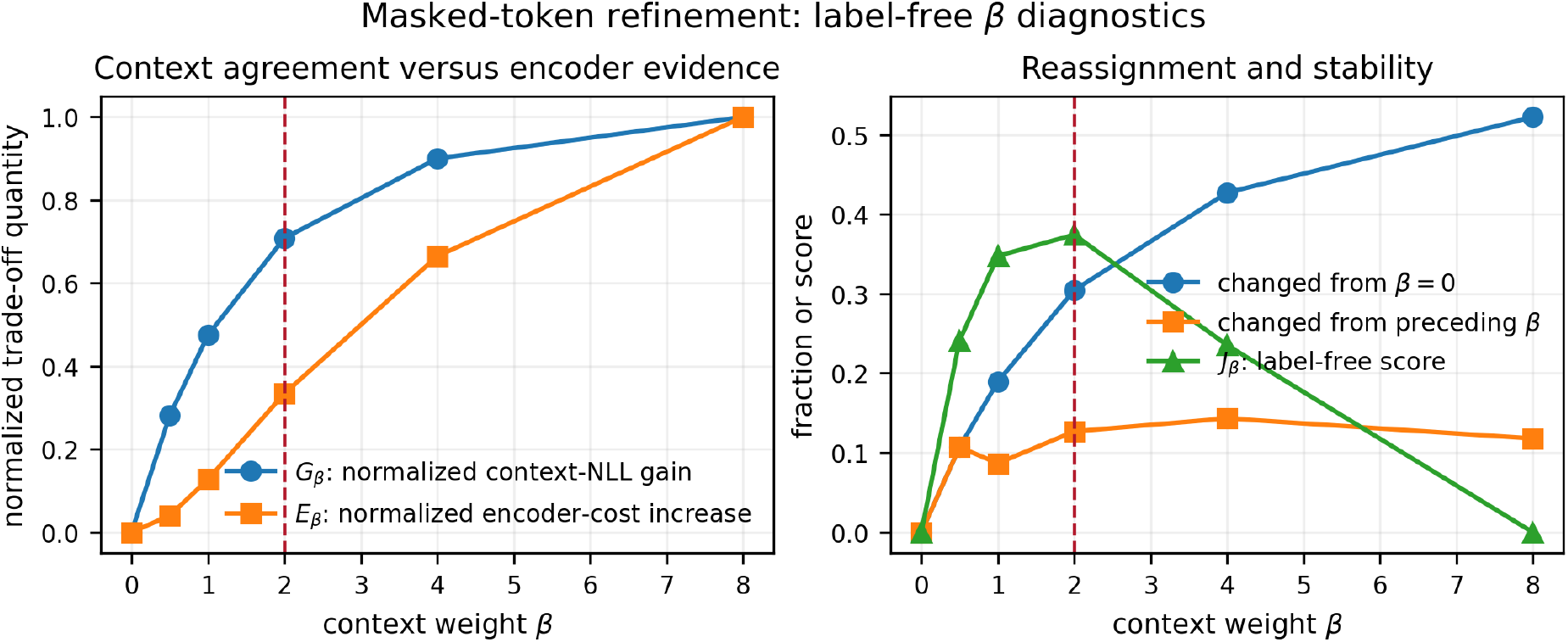
Label-free selection of the contextual weight *β*. The left panel shows normalized context-NLL gain *G*_*β*_ and normalized encoder-cost increase *E*_*β*_. The right panel shows reassigned-site fractions and the label-free score *J*_*β*_ = *G*_*β*_ − *E*_*β*_. The dashed line marks the maximizing candidate, *β* = 2.

The impact of contextual reassignment is illustrated in the progression in Fig. 4. The initial VQ render captures recurring particle structures but is cluttered with weak, diffuse assignments. Refinement using the context tokens concentrates the representation around temporally compatible structures, making the resulting feature states more interpretable. Some true particles are still missing and some noise remains, as expected; contextual compatibility alone does not qualify every occurrence. Therefore, the following stages further isolate the feature of interest from the background and mis-assigned occurrences.

**Figure 4:**
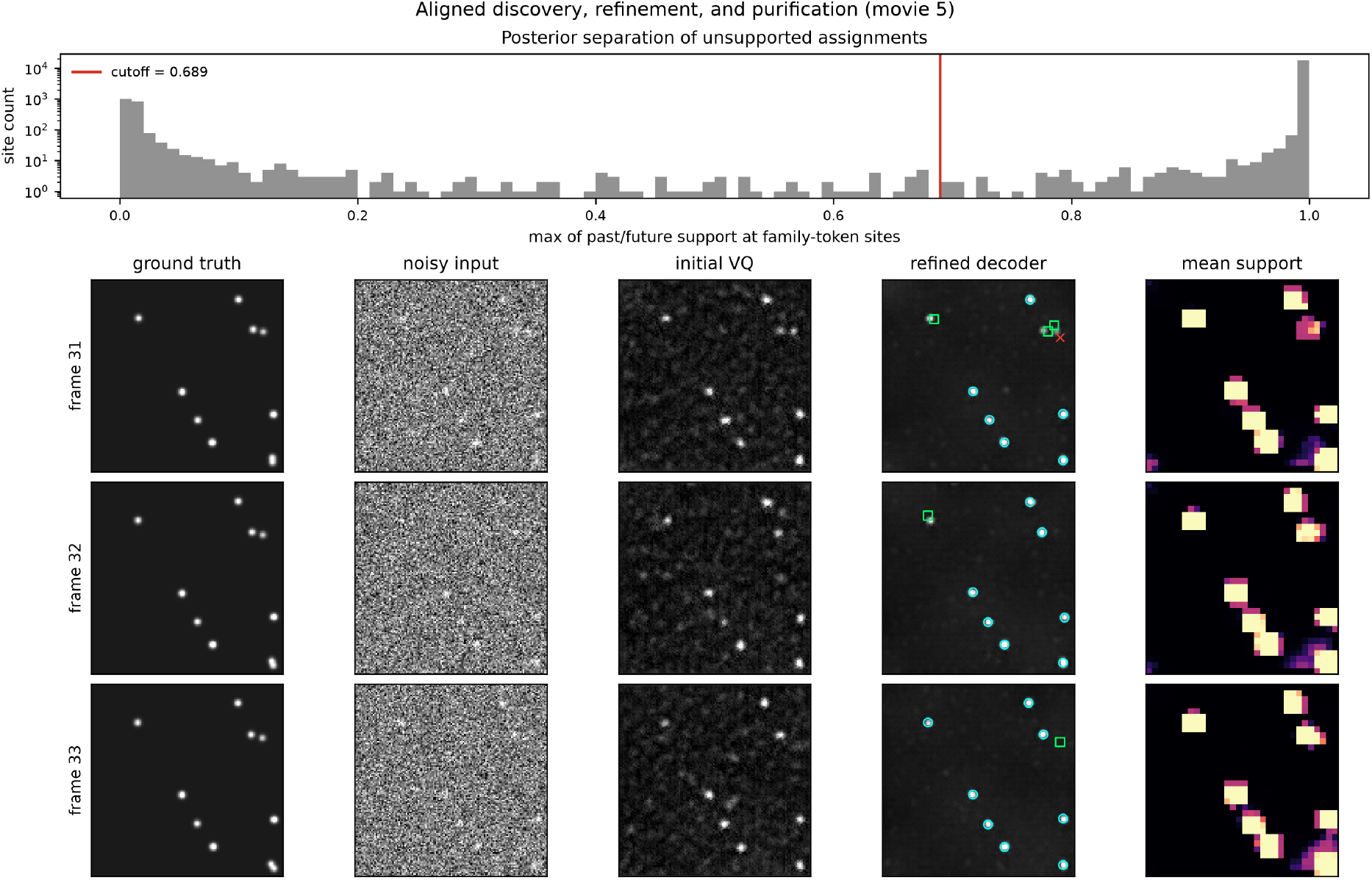
Three consecutive frames from the same generated video. The upper panel gives the distribution of the maximum of past and future support and its 0.95 posterior cutoff. In the refined-decoder row, circles mark components retained by the saved component finder, crosses mark family sites selected for suppression, and squares mark high-support non-family candidates near otherwise missed synthetic particles. The squares indicate potential candidate with high support scores but missed with the featured codes. The final column shows the arithmetic-mean of the past and future branches.

### 3.2 Stage B&C: Feature-Family Definition and Analysis

Using the family-discovery procedure from Sec. 2.6, candidate anchors are identified as local maxima of the joint activity–proposal statistic that exceed the 0.95 activity and 0.90 proposal quantiles. State codes are ranked with the samples in this pool according to Eq. 13. Figure 5 displays the resulting enrichment and normalized patches after decoder refinement. States 5 and 8 were the most representative tokens and thus formed the *feature-family* for subsequent analysis, while state 11 was confirmed as *background*. This step effectively assigned the project-specific label “PSF-like” to these learned states. Note that the averaged patches remain similar to Fig. 2, although more than 30% of the initial assignment are altered (Fig. 4). The improved rendering therefore indicates that contextual refinement primarily corrects feature sites that were initially mis-assigned to noise/background states.

**Figure 5:**
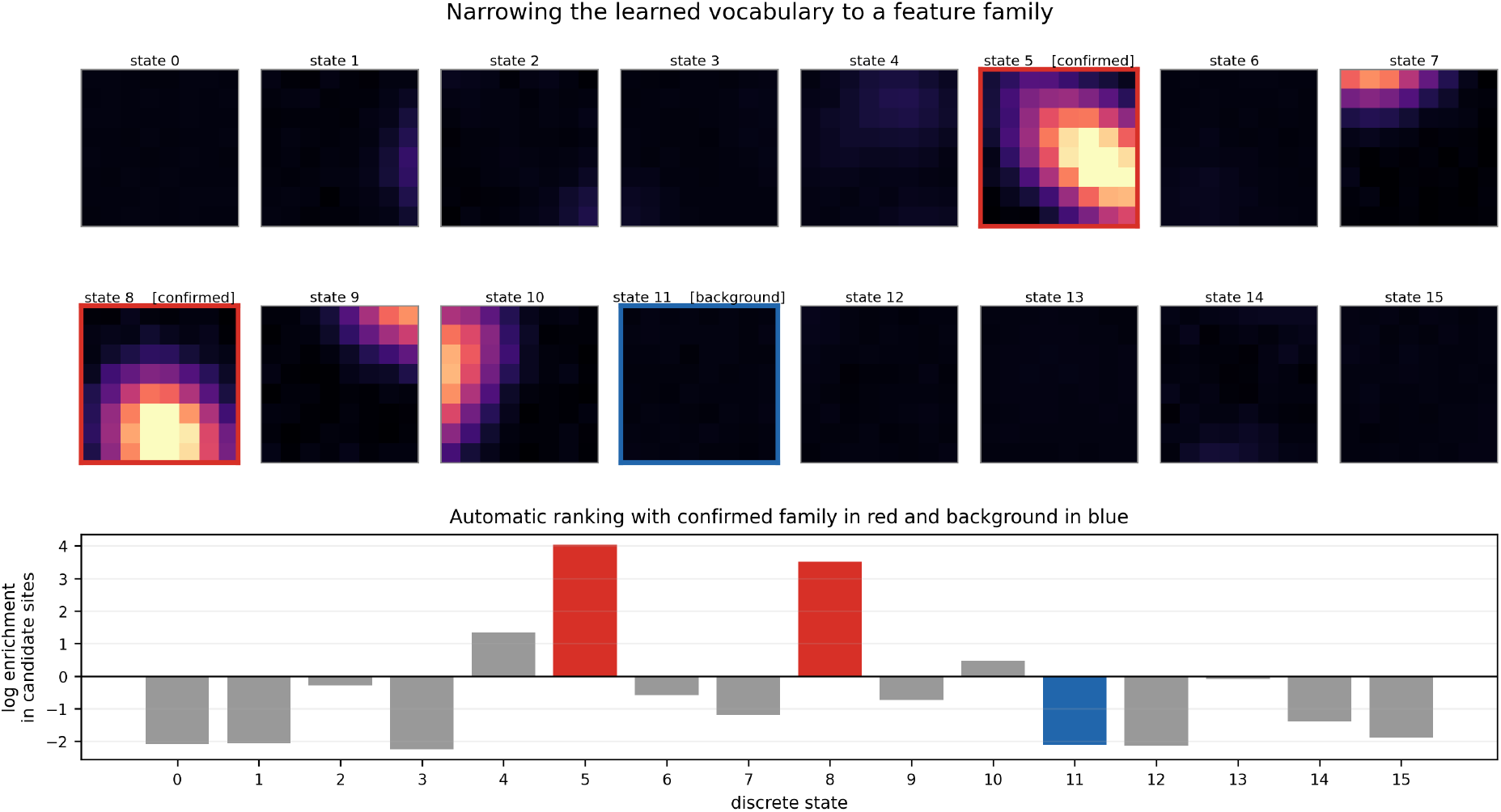
Feature-family selection. The atlas shows normalized patches associated with the refined tokens; the lower panel gives their enrichment at high-activity, temporally consistent candidate sites. States 5 and 8 form the feature-family, and state 11 is the selected background.

The directional support classifiers from Sec. 2.7 are implemented with a temporal radius of three frames and a spatial radius of one latent site. The combined-support distribution yields a 0.95 posterior boundary of 0.689 (Fig. 4, top panel). Of the 20,260 strict-family sites, 2162 (10.7%) fall at or below this threshold and are replaced by the background code per Sec. 2.8. A retrospective check shows that 26 of these removed sites lie within 6 pixels of a true particle coordinate, meaning 1.2% of removals are potentially erroneous. This figure is an upper-bound diagnostic rather than a precise false-removal rate, as a single particle can be represented by multiple neighboring latent sites. Conversely, sites with high support but low rendered intensity or non-feature tokens suggest missed detections—cases where temporal evidence is strong despite a weak or non-feature current-frame representation. These are marked with squares in Fig. 4.

Finally, the learned representation is compared against the clean synthetic reference. Figure 6 shows an average of raw noisy-image patches centered at coordinates from the component finder (sampling 5414 patches from 8311 retained components fully within the frame), compared against 10,843 clean-image patches aligned at known particle positions. Before comparison, each pixel-wise average was background-subtracted using the median of its boundary pixels and normalized by its maximum. The similarity between these radial profiles demonstrates that the retained components recover the characteristic PSF morphology from raw data.

**Figure 6:**
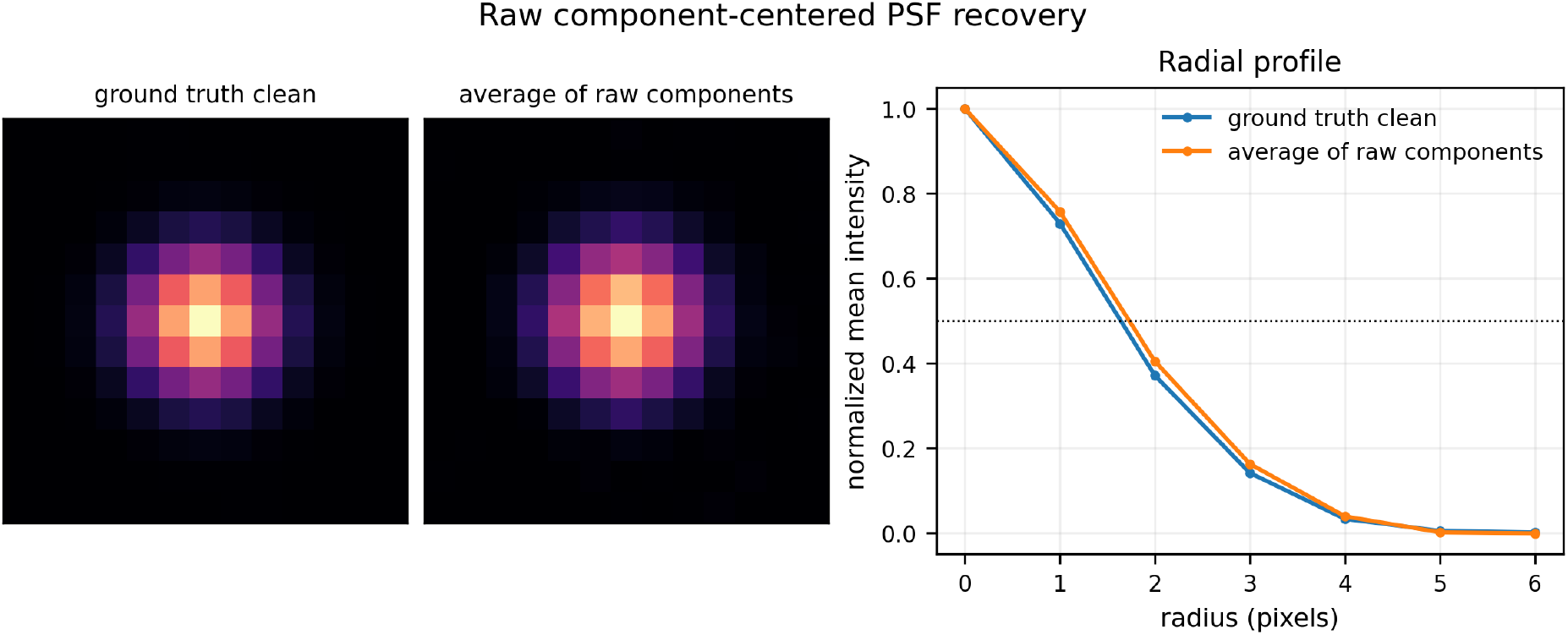
Raw component-centered PSF recovery in the synthetic collection. The recovered average uses raw patches centered at retained component coordinates after suppression; the clean reference are aligned at known particle coordinates. Both averages are boundary-median subtracted and maximum normalized. Ground-truth coordinates do not enter component selection or recovered-patch alignment.

## 4 Test Cases

### 4.1 Low-SNR Particle-Tracking Benchmark

The particle-tracking benchmark from Chenouard et al. [16] is used, which offers simulation ground truth across a range of motion patterns, densities,, and signal strengths. Following its Poisson noise model for photon counts, the operational signal-to-noise ratio is defined as 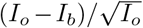, where *I*_*o*_ and *I*_*b*_ represent peak particle and mean background intensities. In contrast to the synthetic data in Sec. 3, the PSFs in this benchmark have a constant intensity. Since conventional tracker performance typically drops sharply below SNR = 4, the framework is tested on the more demanding SNR = 2 videos, covering all three provided densities and two motion scenarios. Here, Vesicles exhibit two-dimensional Brownian motion, while Receptors alternate between Brownian and randomly oriented directed motion, including instances where particles appear or vanish. Receptor is a fast scenario relative to Vesicle and the dataset in Sec. 3. The model operates without any knowledge of these motion laws or scenario labels.

A separate representation for each scenario is learned across the three density levels. Both runs employ *u* = 8 pixels, *K* = 16, a temporal radius of *r* = 2, and the geometry specified in Eq. 1. The label-free context sweep yields *β* = 1 for the Vesicle data and *β* = 2 for the Receptor data. Directional support is computed with a spatial radius of one latent site and a temporal radius of three frames. Feature-family and support boundaries are derived directly from the noisy data. The resulting states are then identified via automated ranking and manually confirmed as “particle-like,” independently of any particle template or motion model.

Two different component definitions are used for these tests. For the Vesicle scenario, a simple one-feature-site component is used whose center must belong to the confirmed feature set. For the Receptor scenario, the structured-component definition from Sec. 2.9 is applied, where the auto-selected feature family serves as the central set, ten context codes are identified, and a component consists of a central-set site paired with at least two context sites within its radius-one Manhattan cross.

Table 1 details frame-wise component detection instead of full trajectory linking. Components are matched one-to-one with true particles using a 6-pixel radius, which roughly matches the simulation’s Rayleigh disk size. To avoid boundary artifacts, the final rendered frame is excluded and a spatial margin of 8 pixels. Precision is defined as the fraction of matched detections, while recall is the fraction of matched true particles. Localization error is calculated for matched components only, and adjacent-pair recovery tracks how often the same ground-truth particle is detected across two consecutive frames. The Jaccard Similarity Coefficient provides a single measure that unifies precision and recall.

**Table 1:** Component detection on the Chenouard SNR = 2 videos. Values use the boundary-excluded scope and a 6-pixel one-to-one matching radius. Error is median frame-localization error among matched components. JSC stands for Jaccard Similarity Coefficient.

| Scenario | Density | Precision | Recall | Pair recovery | Error (px) | JSC |
| --- | --- | --- | --- | --- | --- | --- |
| Vesicle | low | 92.4% | 74.5% | 63.6% | 2.39 | 70.2% |
| Vesicle | mid | 92.8% | 50.9% | 32.0% | 2.35 | 49.0% |
| Vesicle | high | 94.5% | 32.2% | 13.8% | 2.32 | 31.6% |
| Receptor | low | 87.0% | 71.6% | 59.5% | 1.20 | 64.7% |
| Receptor | mid | 91.6% | 50.8% | 35.5% | 1.37 | 48.5% |
| Receptor | high | 94.4% | 41.7% | 26.6% | 1.47 | 40.7% |

For low and medium densities, precision fell between 87.0% and 92.8%, with recall from 50.8% to 74.5%. Precision climbed above 94% at high density, but recall and consecutive-frame recovery both declined, likely because particle proximity led to merges, omissions, or rejections. In the Receptor run, for example, recall dropped from 71.6% to 41.7% and adjacent-pair recovery fell from 59.5% to 26.6% as density rose. The current training mixes all three densities. Its impact will be discussed in Sec. 5. For context, the Jaccard Similarity Coefficient (JSC) is comparable to those in Chenouard et al. ([16] Supplementary Table 2), where it would potentially rank first or second. This is a contextual rather than a direct comparison, however, as the present results focus on frame-wise detection while the cited values describe full tracking. Overall, these tests show that a selective low-SNR particle population can be recovered without needing prior noise or motion models.

High-density data additionally produce learned states that encode contact context. One of the enriched central states in the Receptor scenario, code 4, displays PSF contact along the left diagonal. Figure 7 shows its refined average patch and three examples from the high-density video. Each listed structured component contains exactly one code-4 site and at least two central-set sites, using color to distinguish the contact-associated role of code 4. Ground-truth coordinates were only used post-inference to identify and annotate these close-particle examples.

**Figure 7:**
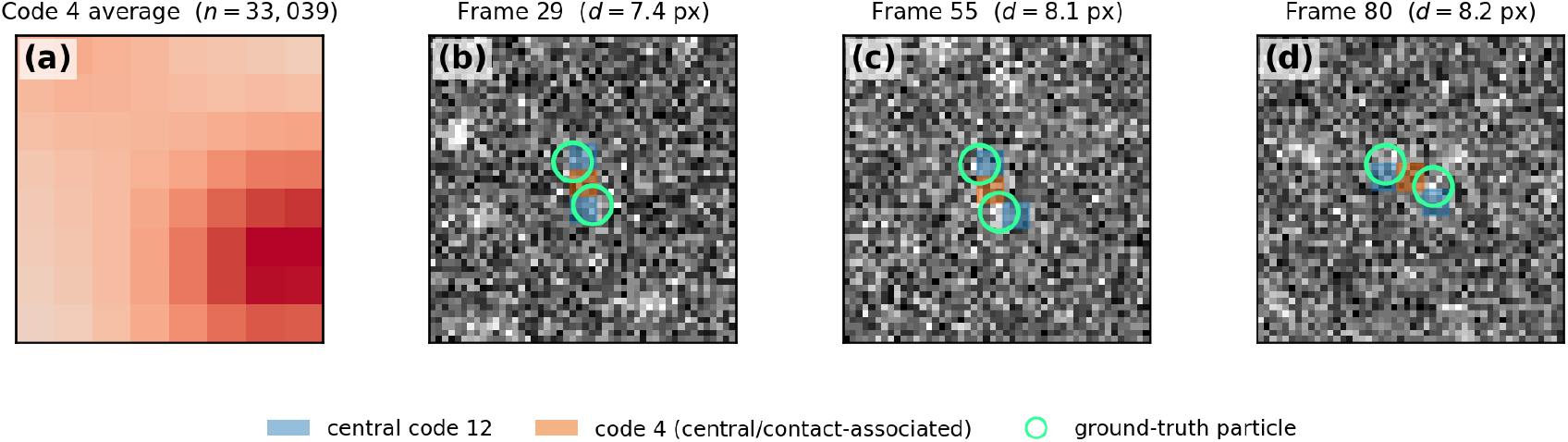
Contact-associated code 4 in the high-density Receptor video. (a) Refined normalized average patch for code 4. (b)–(d) Raw-image crops centered on three retained structured components, each containing at least two central-set sites and exactly one code-4 site. Each marked ground-truth particle lies near a distinct central-set site. Blue marks the other central-set sites, while orange marks code 4 in its contact-associated role. Ground truth was used for post hoc example selection and annotation, not for model processing.

### 4.2 Experimental *E. coli* Video

An experimental video of *E. coli* suspension demonstrates the removal of artifacts. The microscopic images are obtained with a resolution of 4 pixel*/*μm. The rod-shaped *E. coli* bacteria have a diameter of 1 μm and a typical length of 3 μm. Depending on their depth relative to the focal plane, the bacteria appear as dark or bright objects against a gray background. The images contain nonuniform illumination and weak line-pattern acquisition artifacts (see Fig. 8a). A *K* = 64 vocabulary is used to capture the variety of bacterial orientations and acquisition states, which are more diverse than the symmetric-PSF cases. The token map is then refined via contextual reassignment (*β* = 8). This run bypasses feature-family selection and support-based purification. Instead, codes representing artifacts are manually selected and suppressed during rendering. This exploratory edit aims to isolate negative artifacts beyond simple noise, rather than defining positive features. The first artifact addressed is uneven illumination (see Fig. 8a).

**Figure 8:**
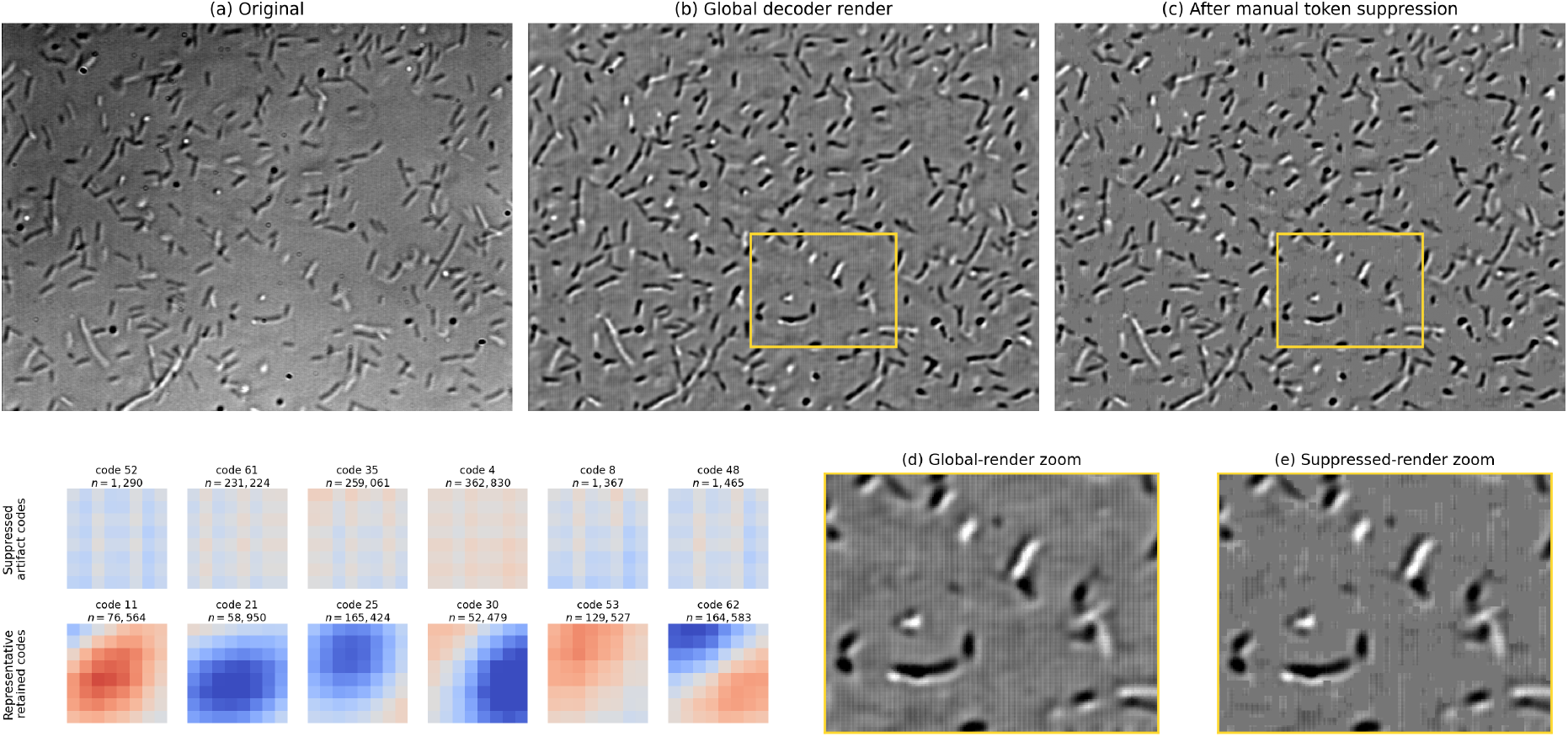
E. coli render-time suppression example. The top row (a)-(c) compares the original frame, the globally scaled refined-decoder render, and the same render after the six manually selected codes were replaced by normalized background during rendering. The lower-left block compactly shows 12 average patches: every suppressed code and six representative retained codes. Yellow boxes mark the common region enlarged at lower right before and after suppression. This manual intervention illustrates codebook interpretability but is not the automated directional-support rule used in the current pipeline.

As described in Sec. 2.2, the refined decoder first maps the token grid to a normalized image *D*_*t*_. Restoring the location *l*_*tij*_ and scale *q*_*tij*_ saved at each latent site gives a local-scale render,

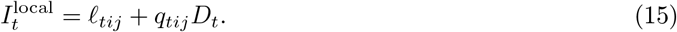

In the current testing video, two distinct sources of unwanted artifacts can therefore appear in the render: the saved site-wise affine field (*l*_*tij*_, *q*_*tij*_), which carries uneven illumination, and the token-dependent image *D*_*t*_, which can carry acquisition artifacts once they acquire their own code identities. The global render removes the first by replacing the per-site field with a single robust location *L* and scale *Q* pooled across the video,

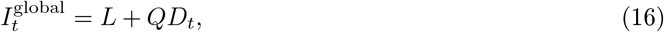

while the token-level step removes the second by suppressing the artifact-associated codes. This two-step removal targets artifacts rather than noise: stochastic noise is not retained as a persistent state and is already attenuated by the temporally masked representation, whereas structured acquisition patterns survive as tokens and require explicit suppression.

Figure 8 compares these different renderings. The global decoder render preserves the bacterial population while removing the local affine field responsible for much of the illumination variation. However, this first step does not eliminate the strip-like fixed pattern noise (FPN); as seen in panels (a) and (b), the model actually learns and enhances this artifact. The subsequent token-level step therefore identifies the six codes associated with the FPN and replaces their footprints in the image grid with *L* (the intensity for normalized zero), while leaving all other codes intact. Comparing panels (b) and (c), and specifically the enlargements in (d) and (e), highlights the impact of these six artifact-associated states. Suppressing them replaces striped or grid-like footprints with the global background without affecting the vocabulary of bacterial appearances. While a Fourier-domain notch filter could remove these stripe frequencies globally, it would also dampen bacterial edges sharing those frequencies. Token-based suppression is local: it only replaces sites assigned to artifact states, leaving the decoder contributions from bacterial states intact. Even though pixel-level restoration is not the goal, this locality is useful to isolate biological features from background artifacts. Since these codes were chosen by inspecting the current run’s codebook, this is a dataset-specific exploratory edit rather than a quantitative evaluation. It shows that acquisition artifacts can manifest as countable, spatially interpretable states that a researcher can explicitly retain, relabel, or exclude.

## 5 Discussion

The demonstration and tests show that temporally masked prediction can concentrate recurrent image structures into a few interpretable states, even without clean targets. This provides a practical means of labeling features within noisy sequences. While codebook learning is self-supervised, the subsequent analysis depends on certain researcher-defined choices (as seen in Fig. 1), including the confirmation of feature families and the definition of components or artifacts. The implications of these choices are discussed below.

### 5.1 Representation Granularity

Vector-quantized representations have proven successful for decomposing cellular images [17, 18, 19]. In this framework, vector quantization serves as both a compression mechanism and an inductive constraint. While a continuous latent space might encode noise-dependent variations across an infinite set of vectors, quantization forces recurrent observations into a finite vocabulary. This allows state occupancy, transitions, and neighborhood relationships to be measured directly. The trade-off is a limit on spatial and appearance resolution; because discretization discards information, sharp or continuously varying structures may be captured by multiple states or approximated by averages. Consequently, the framework focuses on labeling recurrent structures in noisy environments rather than achieving pixel-level reconstruction.

The codebook size *K* determines appearance capacity, while the basic unit *u* governs spatial granularity. Scenes with diverse asymmetric structures, such as the *E. coli* video, typically require more states than those dominated by symmetric features. While *K* = 16 is used for particles and *K* = 64 for bacteria, these results do not define an optimal scaling law for *K*. Instead, *K* should be tuned by monitoring occupancy collapse, state redundancy, and the stability of the selected family, rather than relying solely on reconstruction loss.

The basic unit should roughly match the smallest feature scale of interest. An oversized *u* may blur fine structures and increase the likelihood of merging neighboring objects into a single component, though itself might be an interesting application such as contact detection (see below). Conversely, a *u* that is too small causes a single physical feature to span multiple sites, necessitating a more complex component definition. Equation 1 suggests an even *u* to ensure the latent stride *u/*2 remains an integer; while multiples of four are convenient for implementation, they are not strictly required. Other even values are viable, though odd half-strides can introduce half-pixel shifts in patch-centered visualizations, affecting coordinate interpretation and codebook illustrations more than the underlying token learning.

### 5.2 Density and Acquisition Variation

Object density varies across the datasets in Sec. 3 and Sec. 4.1, affecting both the available training evidence and the ambiguity of the learned states. While sparse videos offer fewer recurrent examples, dense videos introduce more contacts and overlapping appearances. The results of the simulated data in Sec. 4.1 highlight this: while precision remained high at the highest density, recall and adjacent-pair recovery declined. Figure 7 illustrates a potential contact-associated state; under current component definitions, such states may connect and merge two distinct detections. A more refined design could assign these to a specific context role or analyze them as a separate feature family of contacts.

Varying density can also distort the activity distribution, which is critical as activity serves as the anchor for feature definition. To maximize efficiency, the current design uses the 50th quantile to truncate the lower activity distribution (Sec. 2.5), implicitly assuming that half of the pixels are irrelevant background. In extremely dense frames, however, valid features might be misclassified as low-activity even if the VQ model reproduces them accurately. Conversely, in very dilute frames, noise may dominate the distribution and bypass the high-quantile gate during feature-family selection (Sec. 2.6). Mixing high- and low-density data may aggravate this effect: since activity is clipped from below at zero and the median *q*_0.50_ is often zero in low-density frames, the activity distribution develops a zero plateau. This causes the quantiles to allow the entire range to pass the high-quantile gate. This is the most apparent weak point of the current framework. Future iterations should include a distribution check prior to feature-family selection.

Noise levels and illumination artifacts are treated as inherent dataset characteristics rather than nuisance parameters to be ignored. A representation trained in one SNR regime may assign different states to the background of another. While the global rendering mode used for *E. coli* removes the saved site-wise affine field, it cannot erase acquisition artifacts that have been learned into the vocabulary as tokens; these require explicit suppression.

### 5.3 Motion Range and Temporal Interpretation

Since temporal recurrence is the primary source of supervision, motion is measured relative to the basic unit *u* and the temporal radius *r*. If a feature’s displacement over the context interval exceeds the spatial search neighborhood, it can no longer provide reliable local evidence. While expanding this neighborhood allows for faster motion, it also increases the risk of incorporating unrelated sites. Because the framework does not employ a prescribed trajectory to resolve this ambiguity, the selected temporal radius and search extent define an operational range that must be explicitly stated for each dataset.

Using separate past and future classifiers preserves directional asymmetry. Their maximum provides a permissive combined-support score for features that enter, leave, or are temporarily hidden from view, while the separate directional maps record which side supplied the evidence. However, combined support alone cannot establish unique component identity across frames; a nearby but structurally different peak could technically support a central occurrence. The presence of duration-one components in experimental runs is therefore a useful design diagnostic. While possible under the current local support rule, such occurrences are inconsistent with the intended recurrent-feature interpretation and warrant further investigation. Duration constraints and component-level temporal association should thus be evaluated independently of site-wise support.

### 5.4 Future Work

The current framework is intentionally retrospective. Models are trained on sampled crops and frame partitions, then applied to the full dataset under analysis. This transductive approach does not inherently guarantee transferability to new acquisition conditions; such generalization remains outside the scope of this demonstration and requires separate evaluation. Furthermore, the results presented here are not the product of a systematic ablation of human-defined hyperparameters (such as codebook size, basic unit, temporal radius, component definition, or support threshold). Future work should quantify the stability of these choices across diverse acquisition conditions and random initializations.

Finally, several extensions follow naturally from the present framework. First, an explicit orientation variable could be introduced for asymmetric features. Currently, rotated versions of a structure may occupy distinct codes, consuming codebook capacity and splitting a single structural family across multiple entries. A rotation-aware encoder could instead map rotated patterns to a shared content code while estimating the angle as a separate variable, allowing the decoder to reconstruct the patch using both. This could be implemented via rotation-equivariant layers or a discrete set of rotated maps. While unnecessary for symmetric features, this separation would allow for the analysis of orientation distributions and contact geometry in elongated structures without unnecessarily inflating the codebook. A second promising extension is to evolve the support classifier into a full tracker. Because the support map estimates temporal consistency within a bounded search, it provides a natural foundation for trajectory association. While a complete tracker would require additional rules for component-level identity beyond simple pixel intensity, the current framework already enables the study of state changes—such as objects entering or exiting the focal plane—via transition and recurrence statistics, without needing a predefined motion model.

## 6 Conclusion

This work introduces a self-supervised framework for discovering recurrent local structures in noisy image sequences. By combining temporal masking for vocabulary discovery, contextual refinement for evidence reconciliation, and directional support for occurrence qualification, the method extracts discrete features without requiring clean targets. A component finder then groups these sites into operational components for downstream analysis.

Results from synthetic and benchmark tests demonstrate that the framework recovers selective point-feature populations under severe noise, while the *E. coli* example exposes biological structures versus acquisition artifacts. These demonstrations show that the framework supports exploratory analysis of noisy sequences where the relevant features are not defined or assumed in advance. The method is not intended to replace approaches with prior assumptions, such as particle shapes, noise distributions, or motion laws. Instead, the discrete representation complements such methods: retained components can seed a template-based localizer, narrow a tracker’s association search, or initialize a fitting procedure, while the learned states provide a compact descriptor for matching these candidates across frames. Finally, because every step operates on a compact token map rather than raw pixels, training and application are inexpensive enough to be performed per project without a large general pretrained model.

### A Training and Run-Specific Settings

#### Synthetic representation learning

The synthetic demonstration employs 32 videos, *u* = 8, 32 ×32 spatial crops, a temporal radius of two, latent stride four, *K* = 16, and 32-dimensional code vectors. The VQ model is fitted for eight epochs using AdamW (learning rate 2 × 10^−3^, batch size 64) on 4096 deterministically sampled windows (seed 307). Movie and eligible center-frame indices are sampled uniformly. High-variance crops among 16 uniformly drawn spatial candidates are used for twenty percent of windows, while the remainder use one uniform spatial crop, with all crops constrained to lie fully within the frame. The Smooth-L1 reconstruction term is evaluated over the central 8 × 8 pixels. The total objective incorporates the VQ loss—defined as the codebook loss plus 0.25 times the commitment loss—with a weight of 0.25. Median–MAD normalization uses the complete noisy five-frame window, with a MAD floor set to 5% of the global robust scale estimated from the noisy training collection.

#### Contextual reassignment and decoder refinement

In the synthetic demonstration, the masked-token predictor is fitted for four epochs using AdamW (learning rate 2 × 10^−3^, weight decay 10^−4^, batch size 2048), with two complete videos withheld for validation. The saved sweep evaluates *β* ∈ { 0, 0.5, 1, 2, 4, 8 } and selects *β* = 2 according to Eq. 8. While the refined tokens and remaining representation models are frozen, the decoder is refitted for four epochs with AdamW (learning rate 5 × 10^−4^, weight decay 10^−4^, batch size 32).

#### Feature-family and directional-support settings

Candidate-family codes must have an eligible-site frequency of at least 5 × 10^−4^ and at least two occurrences in the gated evidence population. Each directional support model is fitted for six epochs using AdamW (learning rate 2 × 10^−3^, weight decay 10^−4^, batch size 2048). Validation is performed on a contiguous 20% frame block in every movie, with a three-frame guard excluded from training on each side. Background and family-token negatives are sampled at most three and two times the positive count, respectively. The reported mixture cutoff employs a declared lower-component posterior confidence of 0.95.

